# 3D shell asymmetry of Testudines as a potential biomarker for environmental stress

**DOI:** 10.64898/2026.02.04.702268

**Authors:** Merin Joji, Ilya Dziomber, Christy Anna Hipsley

**Affiliations:** Section for Ecology & Evolution, Department of Biology, University of Copenhagen, Copenhagen, Denmark; Konrad Lorenz Institute for Evolution and Cognition Research, Klosterneuburg, Austria; Institute of Plant Sciences, University of Bern, Switzerland; Oeschger Centre for Climate Change Research, University of Bern, Switzerland; Natural History Museum of Denmark, University of Copenhagen, Denmark

**Keywords:** Geometric morphometrics, fluctuating asymmetry, turtle shell, developmental instability, and 3D surface scanning

## Abstract

Turtles and tortoises (Order Testudines) possess a unique bony shell that varies in shape across ecological niches. Previous studies have linked turtle shell abnormalities to the presence of environmental stress, leading to asymmetry in shell shape. Here we present the first large-scale geometric morphometric analysis of shell asymmetry in preserved museum specimens from 92 turtle species, using high-resolution 3D scans and (semi)landmark-based methods. We quantified fluctuating asymmetry (FA) and directional asymmetry (DA) in the whole shell, carapace, and plastron, and tested for ecological and phylogenetic influences on shell shape. Our results reveal significant ecological effects on both symmetric and asymmetric components of shell morphology, with aquatic and marine species exhibiting higher FA than their terrestrial counterparts. The carapace showed higher asymmetry and integration than the plastron, suggesting different developmental constraints. Phylogenetic signal was present but weak, indicating convergence in shell shape among ecologically similar but distantly related species. Partial least squares analysis revealed strong covariation between symmetric and asymmetric components, supporting the shell as an integrated morphological unit. These findings highlight the utility of FA as a non-invasive indicator of developmental instability, with implications for conservation monitoring using preserved and living specimens.

## Background

Turtles and tortoises (Order Testudines) are unique among vertebrates in having a bony shell in which the majority of their body is encased. The shell is composed of the dorsal carapace and ventral plastron, formed by the fused ribs+vertebrae and breast bones, respectively [1]. In the majority of species the shell surface is covered in keratinized scales known as scutes, providing protective armor into which the head and limbs can retreat [2].

Since their origin from a shell-less, terrestrial ancestor around 240 million years ago [3], turtles have diversified into an extraordinary range of environments on every continent except Antarctica [4,5]. Today they inhabit deserts, rainforests, woodlands, lakes, streams and ponds, as well as brackish estuaries and islands, and in the case of sea turtles, the oceans [4,5]. This ecological variation is reflected in the size and shape of their shells and limbs, which vary from the large, streamlined bodies and flippers of marine turtles to the tall, domed shells and club-like legs of tortoises [6–9]. This diversity makes them an ideal group in which to study the effects of environmental stress on morphology, which is urgent as over half of turtle species are currently threatened with extinction—a higher proportion than in any other major vertebrate group [10].

A promising biomarker for detecting environmental stress in diverse taxonomic groups is fluctuating asymmetry (FA) [11,12]. Although many organisms, including turtles, are bilaterally symmetrical (meaning their bodies can be divided into equal, mirror-image left and right halves), random deviations from perfect symmetry can arise due to perturbations in the developmental environment [13]. Resulting asymmetries, while often subtle, may interfere with an individual’s performance or energy allocation to reproduction and development, thus leading to diminished fitness [14,15]. Detrimental effects of FA are likely to be magnified in disturbed environments, where animals are exposed to additional stressors such as pollution, invasive species, and habitat fragmentation. For example, common wall lizards (*Podarcis muralis*) in urban populations display higher FA in head shape than their rural counterparts, indicating disruption to the developmental processes regulating growth in human-altered habitats where temperature and pollution are higher (*Podarcis* having soft-shelled eggs that are sensitive to both [16]). Similarly, *Psammodromus algirus* lizards with higher FA in the hindlimbs experience negative consequences on locomotor performance, with decreased sprint speeds during escape attempts from simulated predators compared to more symmetric individuals [17].

Given that turtles are generally long-lived (up to 190 years for tortoises, or 50 years in captivity for freshwater turtles [18]), experience delayed sexual maturity, and face increasing anthropogenic stress, high levels of FA are likely to constrain their ability to optimally function, survive, and adapt to environmental change [19]. This is particularly relevant for species with temperature dependent sex-determination (e.g., marine turtles), for whom higher temperatures caused by global warming and waste products have the potential to bias sex ratios towards females, thus decreasing population size and genetic diversity [10]. Anthropogenic impacts are already documented across multiple turtle species and habitats [10], including stress-induced disease and mortality in sea turtles [20,21], and phenotypic anomalies in sea turtles, tortoises and freshwater turtles due to teratogens [22–25], entanglement with litter and machinery [26–28], and elevated incubation temperatures [24,29]. Among the observed physical abnormalities, deformities in the turtle shell are frequently reported (Figure 1), indicating developmental instability in the processes underlying skeletal morphogenesis.

**Figure 1.**
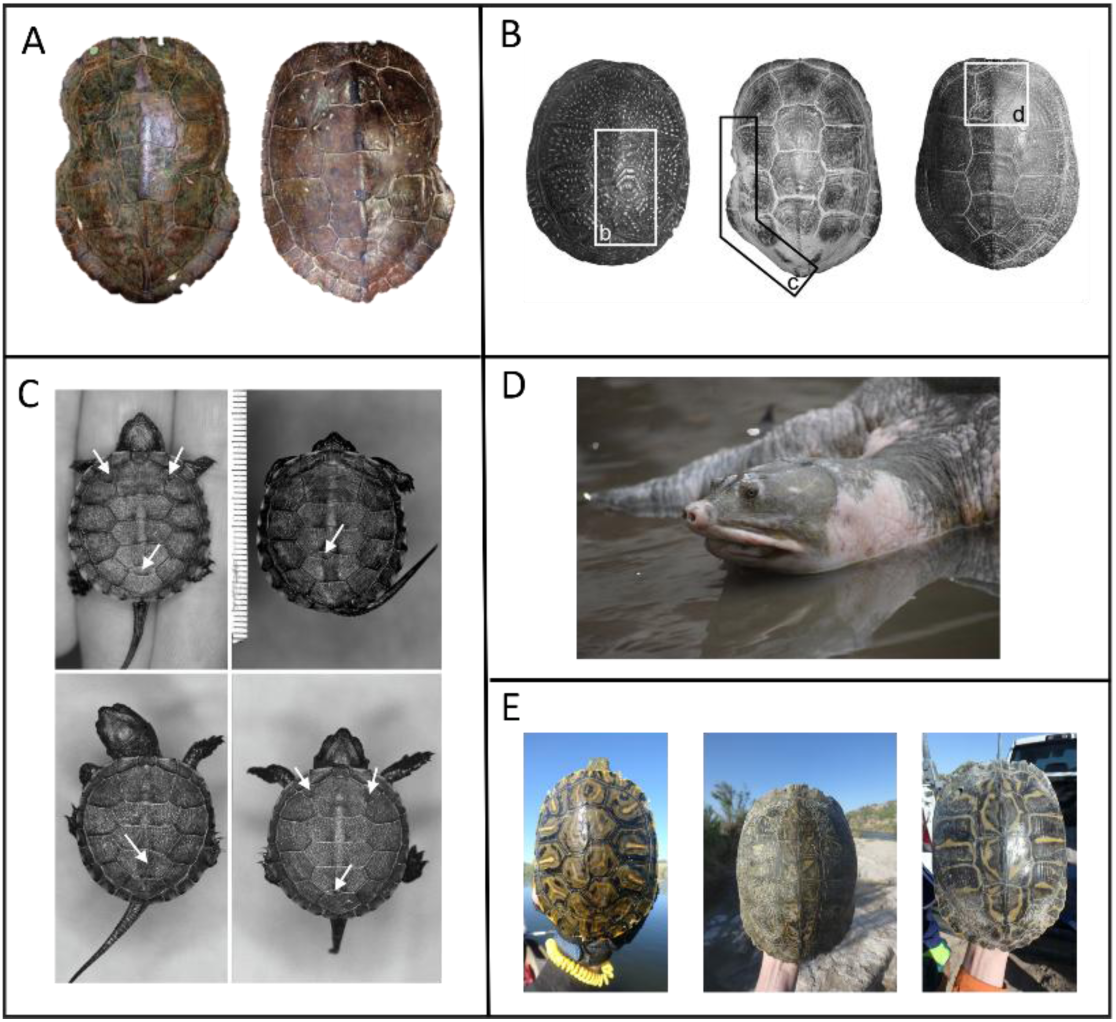
Examples of carapace abnormalities observed in wild turtle populations. A) Deformed marginal scutes of *Graptemys geographica* due to thermal and chemical conditions in nesting substrate [29]. B) Distinct populations of *Emys orbicularis* with abnormal b) accessory scutes, c) shell shape, and d) scute shape, likely influenced by environmental stress and fragmented habitats [30]. C) *Emys orbicularis* hatchling scute anomalies (indicated by white arrows) due to presence of pesticides and heavy metal pollutants [31]. D) Skin discoloration and presence of leaches on the shell of *Nilssonia nigricans* linked to unclean water and lack of basking sites [32]. E) Abnormal vertebral scutes in *Pseudemys gorzugi* turtles living in the Lower Pecos River, New Mexico, USA, caused by poor environmental conditions [33]. Images used with permission from: A) Roy Nagle, B) Marcel Uhrin, C) Adolfo Cordero Rivera, D) Gaurav Bahardiya, and E) Laramie B. Mahan.

Far from dead biological structures like fingernails or antlers, the turtle shell is composed of living tissues that are continuously remodeled and repaired, providing protection and functional adaptations throughout an animal’s lifetime. As the turtle matures, bone growth occurs through osteogenesis along the edges of the shell plates, allowing it to enlarge as the turtle increases in size [34,35]. The layer of keratinous scutes, when present, also grows and is replaced periodically, especially in hatchlings and juveniles [36]. This means that the shell is vulnerable to different developmental stressors at different timepoints, leading to abnormalities or even death. For example, up to 71% of painted turtle (*Chrysemys picta*) clutches exposed to contamination from oil, landfills and sewage were severely deformed, with shell, spine and facial abnormalities often proving fatal [37]. Adult deformities were less severe, although typically involving the feet (missing or jumbled toes), shell (kyphosis or scoliosis), and tail (extra spines at the tip) [37]. While these adults may live to reproduce, developmental damage caused by exposure to environmental contaminants can lead to heritable mutations or epigenetic effects, thus negatively impacting future generations.

In this study, we assess variation in shell asymmetry across Testudines using landmark-based geometric morphometrics (GM) on 3D models of adult specimens. Geometric morphometrics is a more sensitive approach to quantifying FA than linear or meristic measurements [11], as it uses the entire set of landmark coordinates to represent full shape, rather than few, often correlated and size-dependent distances. We furthermore describe variation in shell shape, allometry (the relationship between shape and size) and FA across phylogenetic and ecological groups, to identify potentially vulnerable taxa to current and future environmental stress [12]. Similar methods have been used to evaluate asymmetry in several turtle species [30,38–43], however this paper represents the first large-scale study of FA across all living testudine families, encompassing diverse habitats, diets, behaviors, and reproductive strategies. When combined with field surveys and biomechanical modelling (e.g., finite element analysis [8,44]) to identify the potential sources and consequences of FA on living animals, this approach has the potential to identify populations at risk and to pinpoint healthy individuals for captive breeding or translocation.

## Methods

### 3D model acquisition and landmarking

We analyzed 127 specimens from 92 species of Testudines, representing all 14 extant families (Appendix 1). Sixty-nine surface scan 3D models and their corresponding landmarks were taken from Dziomber et al. [6]. Fifty-eight additional 3D models were acquired from various sources: four from X-ray computed tomography scans provided by Tristan Stayton, one from the Natural History Museum of London, 19 from the online repositories MorphoSource and Sketchfab, and 33 from dry preserved specimens at the Natural History Museum of Denmark (ZMUC). The latter were scanned using an Artec Space Spider (Artec 3D, Luxembourg), which uses structured light and an inbuilt camera to capture the shape and surface (texture, color) of the turtle shell in 3D, creating a high-fidelity model with 0.1 mm pixel resolution. Ecological categories were taken from Dziomber et al. [6], which assigned habitat preferences to species based on forelimb webbing following Foth et al. [45]: not webbed = terrestrial; poorly webbed and fully webbed = semiaquatic; extensive webbing = aquatic; flippers = marine. For the 8 sampled species not included in Foth et al. [45] (see Appendix 1), we assigned ecologies according to the Reptile Database (http://www.reptile-database.org), accessed 15 March 2025.

For the newly acquired specimens, turtle shell meshes were imported into Checkpoint Stratovan software [46] (v 2024.11.14) in .ply format for 3D landmarking. Landmark placement followed the ‘SET1’ configuration of Dziomber et al. [6], which included 10 anatomical landmarks and 12 curves, the latter of which we resampled to 348 semi-landmarks (Figure 2). We hereafter refer to this landmark dataset as the dataset 1. To differentiate between FA and variation in left-right landmark pairs due to measurement error (i.e., inter- or intra-observer differences in landmark placement), all 127 specimens were landmarked again in the same software (hereafter referred to as the dataset 2). These replicated datasets were later used to estimate FA.

**Figure 2.**
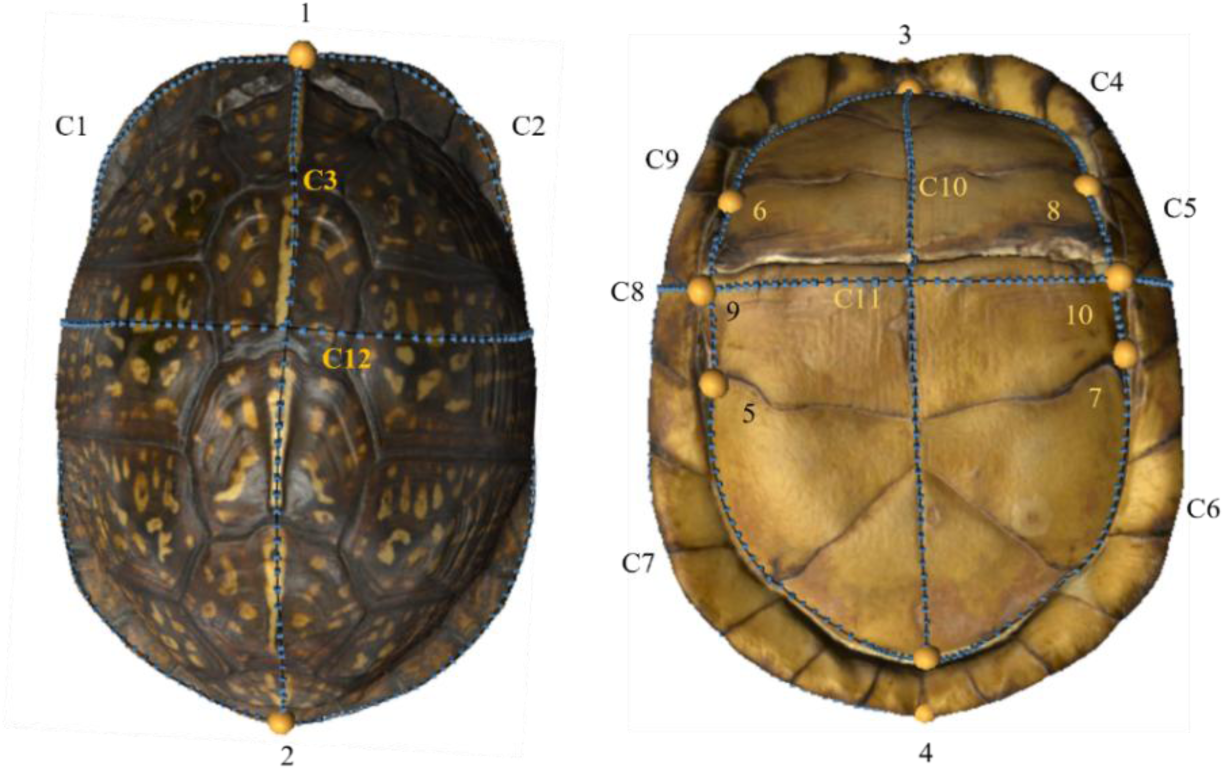
Landmark configuration applied in this study (see Dziomber et al. [6] for details), shown on A) carapace and B) plastron of *Terrapene carolina* (ZMUC R25184). Ten fixed anatomical landmarks (yellow dots) are connected by 11 semi-landmark curves (C1-C11; blue dots). The number of semi-landmarks digitized per curve were: C1 & C2 = 51 each, C3 = 26, C4 = 19, C5 = 15, C6 & C7 = 21, C8 = 15, C9 & C10 = 19, C11 = 25, and C12 = 47.

### Phylogenetic tree for comparative analyses

Related species are statistically non-independent due to shared ancestry, necessitating phylogenetic comparative methods when analyzing their trait variation [47]. We therefore generated a phylogenetic tree in Mesquite v 3.70 [48] based on the maximum clade credibility tree of Thomson et al. [5], pruned to match our dataset. That molecular phylogeny was based on 15 nuclear DNA markers for 279 extant turtle species, with divergence times calibrated by 22 vetted fossils [49]. Some species names were modified to match our sample, while others were manually added at the midpoint of their sister branch according to the following evidence:

1. *Rafetus sp.* was renamed to *Rafetus euphraticus* to align with our sampled specimen [5].
2. *Lissemys ceylonensis* was added as a sister species to *Lissemys punctata* based on morphological and mtDNA evidence [50].
3. *Homopus areolatus* was renamed to *Homopus femoralis,* supported by phylogenetic analysis which recovered them as sister taxa [51].
4. *Psammobates oculifer* was renamed as *Psammobates tentorius*, supported by molecular analysis which also placed *P. tentorius* as sister to *Stigmochelys pardalis* [52].
5. *Trachemys ornata* was added as sister species to *Trachemys scripta* by editing the name of the closely related *Trachemys venusta* [53].
6. *Melanochelys tricarinata* was added as sister species to *Melanochelys trijuga* [54].
7. *Deirochelys chrysea* was replaced with *Deirochelys reticularia*, as it is now considered a subspecies of the latter (*D. reticularia chrysea)* [55].
8. *Emys marmorata* was replaced with *Actinemys marmorata*, supported by recent taxonomic revisions [56].
9. *Lepidochelys kempii* was placed as sister species to *Lepidochelys olivacea,* based on phylogenetic analysis [54].
10. *Natator depressus* was added as sister species to *Chelonia mydas,* as recovered by phylogenetic analysis [54].
11. *Mesoclemmys zuliae* was replaced with the closely related *Mesoclemmys dahli* [57].
12. *Chelodina mccordi* was replaced with *Chelodina oblonga* [54].
13. *Emys blandingii* was renamed to *Emydoidea blandingii*, based on recent taxonomic revisions in the family Emydidae [56].

The resulting tree with labelled species ecologies is shown in Figure 3.

**Figure 3.**
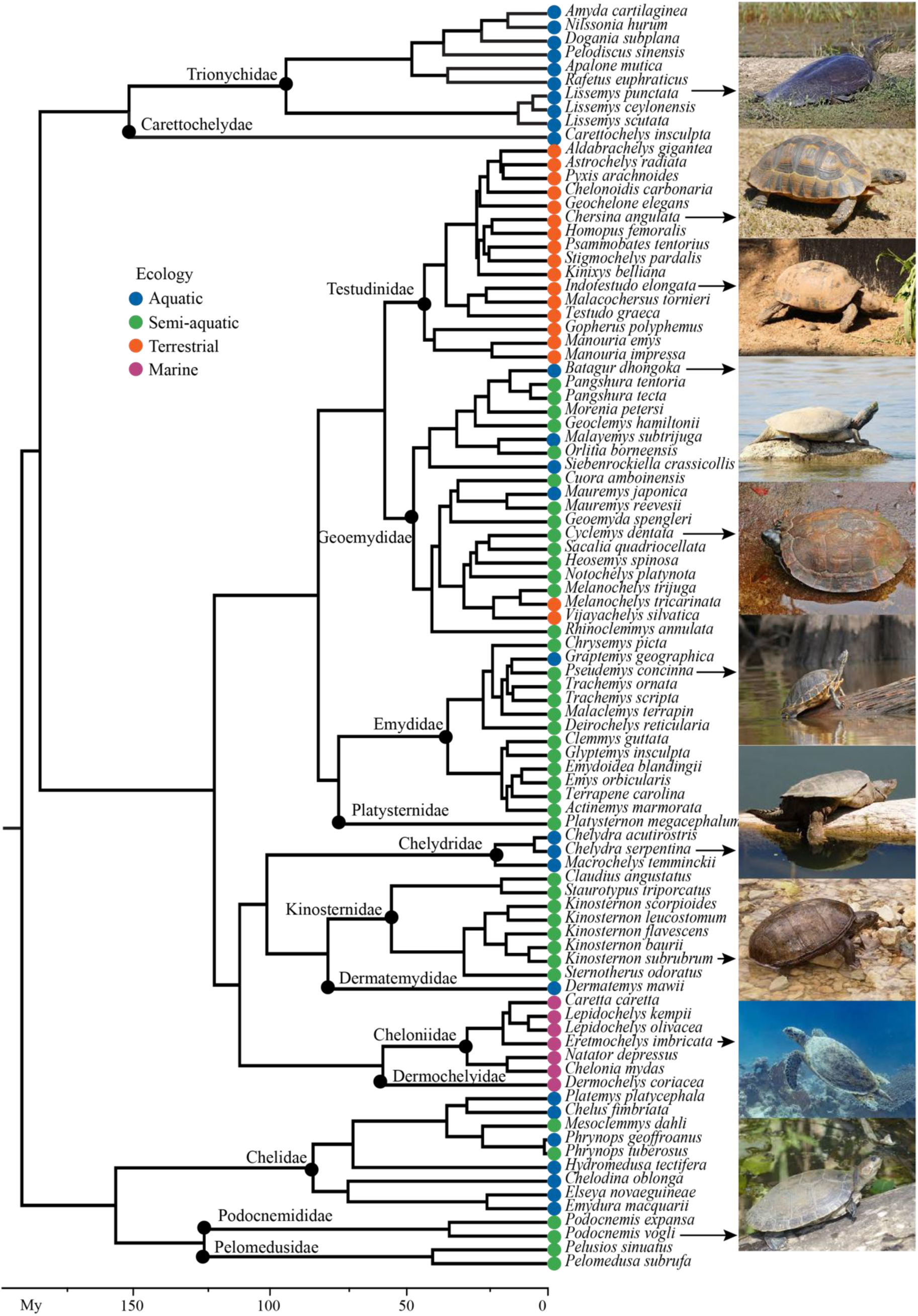
Phylogenetic tree used in geometric morphometric analyses of Testudines, with families indicated by black dots. Divergence dates in millions of years (My) before present are shown along the x-axis. Colored balls at the tips of branches indicate species ecologies, with images representing select species in their natural habitats. Images are shown under Creative Commons licenses CC BY-SA, CC BY-SA 2.0, CC BY-SA 3.0, and CC BY-SA 4.0.

### Geometric morphometric analyses

Analyses were carried out in the R statistical environment v. 4.4.1 [58], using the geomorph package (v 4.0.10) [59] with functions listed in italics below. *Digit.curves* was used to specify the number of semi-landmarks along each curve (see Figure 2 for landmark numbers). Because Procrustes analysis requires the same number of landmarks for all specimens, we used *estimate.missing* to estimate the coordinates of missing landmarks based on thin-plate spine interpolation (method= “TPS”) [6]. This was applied to two turtles with three missing landmarks each, and one turtle with six missing landmarks. A generalized Procrustes analysis was performed using *gpagen*, thereby scaling, rotating, and translating landmark configurations to a single coordinate system. The default option of minimizing bending energy (ProcD = FALSE) was chosen to slide semi-landmarks along their curves by adjusting their positions along the tangent vectors until shape dissimilarities among specimens are minimized [60]. This step generates a set of Procrustes-aligned coordinates, which were used in the following tests. For all analyses, 10000 permutations were applied to estimate statistical significance.

To explore allometric, ecological, and phylogenetic patterns of shape variation, we took species averages of the two landmark (dataset 1 + dataset 2) and incorporated the above tree in phylogenetic analyses. Linear regression of Procrustes shape coordinates on size and ecology was performed using *procD.lm*, in which size was represented by log-transformed centroid size. In landmark-based GM, centroid size is defined as the square root of the sum of squared distances of landmarks from the center of their configuration [61], often used as a proxy for biological size. Phylogenetic generalized least squares (*procD.pgls*; also known as phylogenetic ANOVA [47]) was used to assess evolutionary allometry among species, and to test the relationship between shell shape, size and ecology while considering shared ancestry. When significant effects were detected, *pairwise* from the RRPP package (v. 2.1.2) [62] was used to assess the magnitude of shape variation among ecological groups, by slope vectors (test.type = “VC”) [63,64]. Eighteen of the 3D models from MorphoSource and Sketchfab had incorrect scaling (see Appendix 2); these were therefore removed from allometric analyses.

The phylogenetic signal of shell shape was estimated using *physignal.eigen* – a multivariate version of Blomberg et al.’s [65] *K*-statistic that evaluates the degree of phylogenetic structure in Procrustes shape variables compared to that expected under a Brownian Motion (BM) model of evolution [66,67]. Here, we used species scores from the first 20 principal components of the below morphospace (representing 95% of the total shape variation) as input variables to remove redundant dimensions, allowing the determinant and trace of **K** – the matrix generalization of Blomberg’s *K* – to be computed [66]. This function calculates two additional metrics: *KA* is the arithmetic mean of the eigenvalues of the phylogenetic signal matrix **K** (i.e, the trace of **K** divided by the number of variables), reflecting the average signal across trait dimensions, while *KG* is the geometric mean of these eigenvalues, representing the ratio of generalized variances between observed and phylogenetically corrected data (the determinant of **K**) to the power of the number of variables. Here we report the determinant and the trace of **K**, noted as detK and traceK, respectively, as statistics tests are performed on these values and not directly on *KA* and *KG* in the geomorph package. For comparison, we also report K_*mult*_, the trace ratio of observed and corrected sum of squares and cross-products matrices, which provides a global measure of phylogenetic signal but is less sensitive to signal heterogeneity across shape space dimensions [66–68]. To visualize evolutionary patterns of Testudines shell shape, the phylogenetic tree in Figure 3 was projected into morphospace of the 92 species using *gm.prcomp*, with the position of internal nodes determined by ancestral state reconstruction [69].

### Shell asymmetry estimation

We used *bilat.symmetry* to quantify asymmetry in all 127 models from the dataset 1 and 2 [16]. The function works by reflecting the landmarks from one side of the shell midline onto the opposite side, enabling the decomposition of shape variation into symmetric and asymmetric components [16,70,71]. The symmetric component represents the shared shape features across both sides, while the asymmetric component captures the differences between them [70,71]. This allowed us to distinguish between FA, which may reflect developmental perturbations, and directional asymmetry (DA), which refers to consistent left-right differences across individuals (e.g., one side being systematically larger) [70,71]. The model also accounts for measurement error through the interaction of individual, side, and replicate [70,71].

In addition to whole shell asymmetry, FA values were estimated for the carapace and plastron landmark configurations separately using the method described above. The raw coordinates were partitioned into their respective portions while discarding C11 and C12 transverse curves (see Figure 2), as they contained landmarks on the shell articulation (hinge) as opposed to dorsal and ventral shells only. Because the generalized Procrustes fit standardizes each configuration to unit size based on different numbers of landmarks (whole shell = 358, carapace = 132, plastron = 152), the resulting FA values cannot be directly compared between datasets [72]. To visualize asymmetry shape patterns among species, the asymmetric shape component was plotted as arrow plots for each configuration using *ggplot2* in R [73]. We used the unsigned asymmetry index, calculated as the square root of sums of squared deviations of left-right landmark pairs [16], to evaluate individual FA variation in the whole shell, carapace and plastron, with statistical differences among ecological groups determined by ANOVA. Finally, we assessed covariation between the symmetric and asymmetric components of shape variation for the carapace and plastron using two-block partial least squares (PLS) analysis (*two.b.pls*) [74].

## Results

There was a weak but significant correlation between shell shape and log-transformed centroid size among individuals (R^2^ = 0.024, *p* = <0.0001) and species, the latter both with and without phylogenetic correction (R² = 0.09, *p* = <0.0001; phylogenetic ANOVA: R² = 0.056, *p* = 0.002). Allometric trajectories varied among ecological groups (R^2^ = 0.123, *p* = <0.0001), with large aquatic and marine species tending to diverge in shape from semi-aquatic species of similar size (Figure S1). Less than one-fourth of the variation in shell shape was predicted by ecology (R^2^ = 0.197, *p* < 0.0001; phylogenetic ANOVA: R^2^ = 0.172, *p* = 0.006), with the greatest differences observed among aquatic and terrestrial turtles, followed by marine species vs. all other groups (Table 1). The smallest differences in mean shape were between semi-aquatic and both aquatic and terrestrial taxa (Table 1). Shell shape displayed weak phylogenetic signal, indicating that species resemble each other less than expected under a BM model of trait evolution (K_*mult*_=0.0216, Z=4.482; traceK**=**6.414, Z=4.65; detK=4.003 x 10^-15^, Z=9.103; *p* for all tests ≤ 0.001).

**Table 1.**
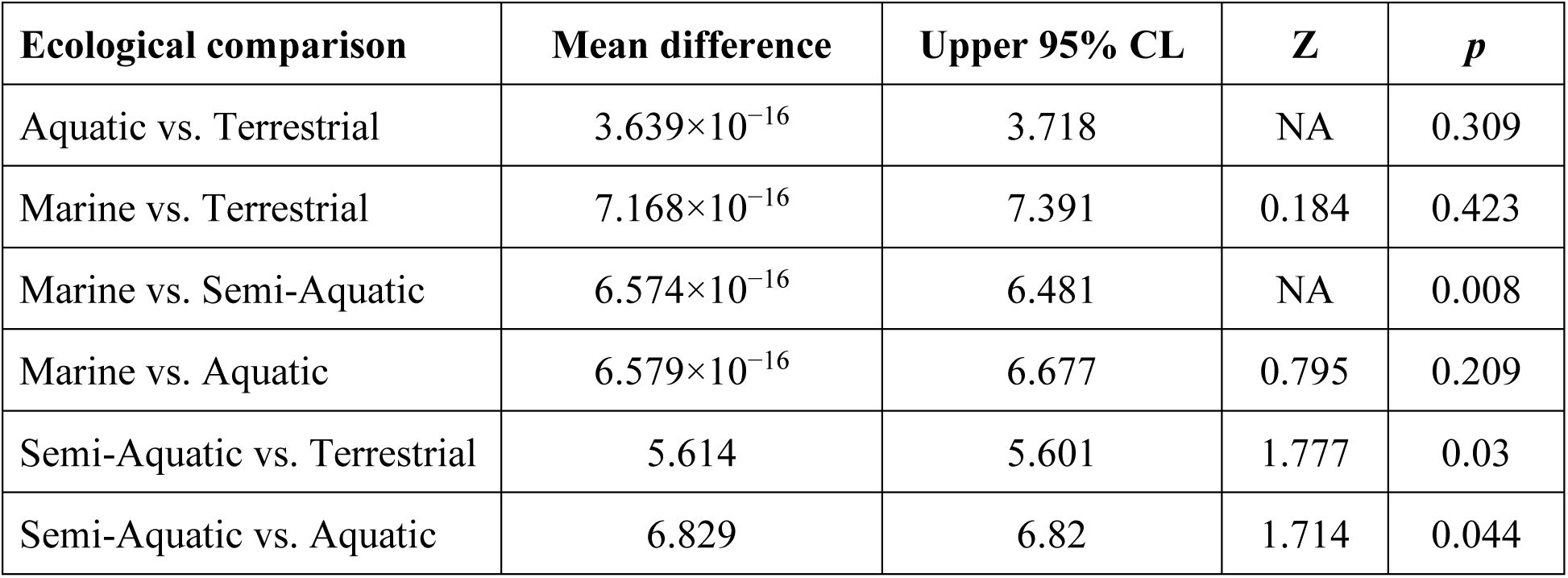
Pairwise comparisons of turtle shell shapes among ecological groups, in order of most to least different. Mean difference represents the Euclidean distance between group mean shapes, listed with the upper 95% confidence internal (CL).

Species were widely dispersed in phylomorphospace, with loose ecological groupings (Figure 4). The first two PCs accounted for nearly half (49%) of the total shape variation (Figure 4; the remaining PC axes described 9% or less). PC1 (35%) captures a gradient from broad, flattened shells, represented by the smooth softshell turtle (*Amyda cartilaginea*), which occupies the low end of PC1, to elongated, domed shells such as the radiated tortoise (*Astrochelys radiata*), which lies at the high end of PC1. PC2 (14%) represents variation in shell height and lateral expansion, separating highly flattened aquatic and marine turtles along the low end of PC2 from more domed forms such as the common snapping turtle (*Chelydra acutirostris*) on the positive end.

**Figure 4.**
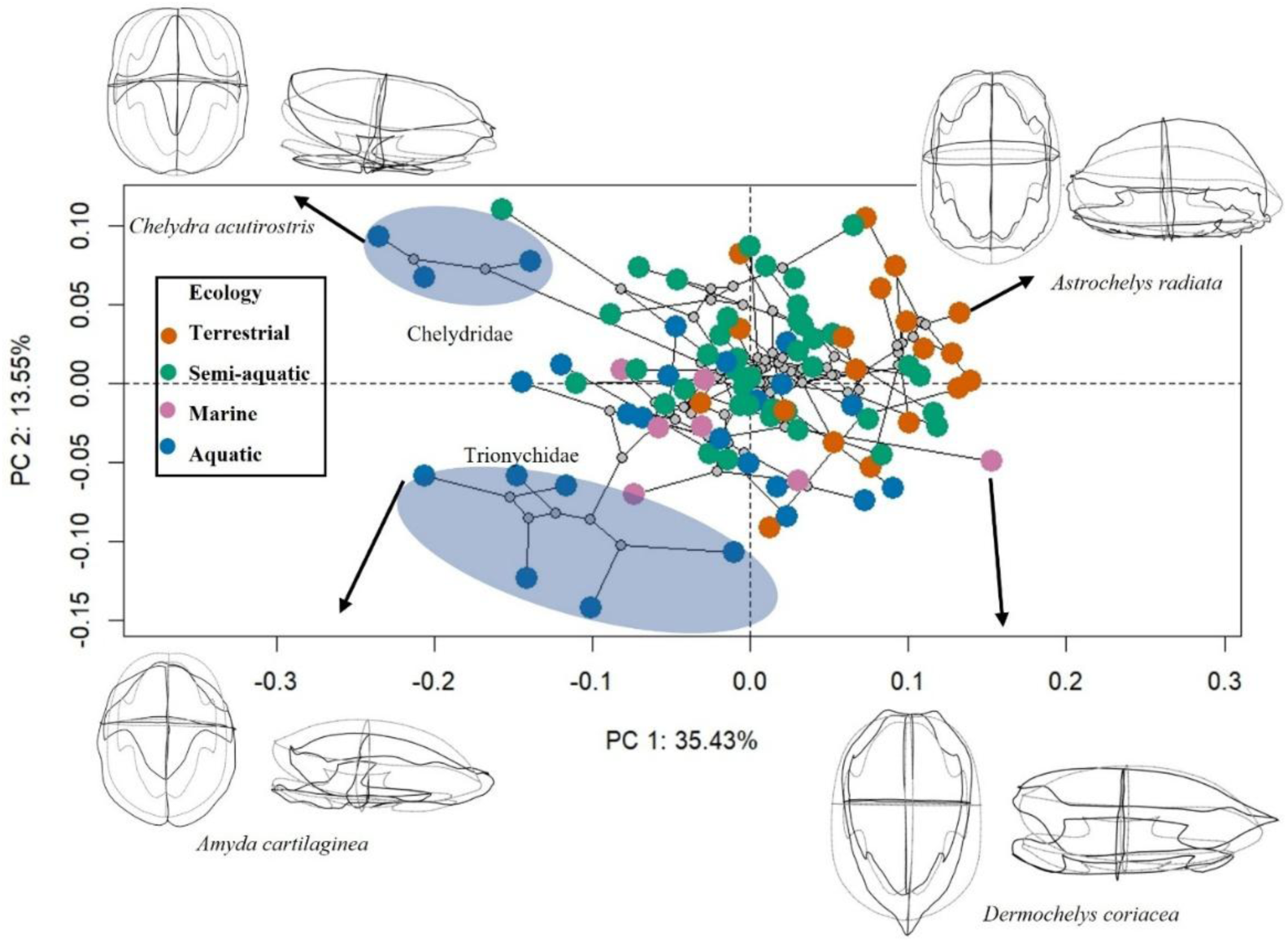
Phylomorphospace of 92 turtle species based on principal component analysis of whole shell 3D landmarks. Species are colored according to ecology. Wireframe diagrams illustrate the mean shell shape (grey) and species-specific variation (black) for selected taxa at phenotypic extremes. Grey dots connecting branches represent estimated ancestral states. Two families forming distinct clusters are marked with blue circles.

Terrestrial taxa tend to have positive PC1 values in opposition to other taxa having negative values. Semi-aquatic species were more broadly dispersed, occupying intermediate regions. Notably, the leatherback sea turtle (*Dermochelys coriacea*), the only living member of its family Dermochelyidae, is placed distantly from its closest relatives the Cheloniidae, containing the remaining six species of marine turtles. This reflects its distinctive tear drop-shaped shell morphology, which contrasts with the more oval to elliptical shell shapes typical of other marine turtles.

### Asymmetry in turtle shell shape

Procrustes ANOVA revealed distinct patterns of asymmetry across the whole shell, carapace, and plastron (Table 2). Side effects were highly significant for the whole shell (accounting for 83% of total variation) indicating strong DA, whereas for the carapace and plastron, side effects were minimal (∼1%). In contrast, highly significant individual effects dominated for the carapace and plastron, reflecting substantial shape variation among individuals rather than between sides. For whole shell, individual effects were weaker and nonsignificant. Fluctuating asymmetry (FA), represented by the individual-by-side interaction, was small across all regions, and significant for the whole shell and carapace, but not for the plastron. Measurement error was generally low, accounting for 1% of the total variation in the whole shell, 3.3% in the carapace, and 8.6% in the plastron.

**Table 2.** Procrustes ANOVA results of bilateral symmetry in the turtle carapace and plastron.

| | df | SS | MS | $R^2$ | $F$ | $Z$ | $p$ |
| --- | --- | --- | --- | --- | --- | --- | --- |
| **Total shell** |  |  |  |  |  |  |  |
| Ind | 126 | 8.401 | 0.067 | 0.14 | 15.529 | -8.809 | 1 |
| Side | 1 | 50.08 | 50.08 | 0.835 | 11664.03 | 5.962 | <0.0001 |
| Ind:side | 126 | 0.541 | 0.004 | 0.009 | 1.2156 | 3.592 | 0.0002 |
| Ind: side: replicate | 254 | 0.897 | 0.004 | 0.01 |  |  |  |
| Total | 507 | 59.920 |  |  |  |  |  |
| **Carapace** |  |  |  |  |  |  |  |
| Ind | 126 | 4.562 | 0.036 | 0.946 | 51.815 | 16.581 | <0.0001 |
| Side | 1 | 0.066 | 0.006 | 0.001 | 9.4718 | 4.8714 | <0.0001 |
| Ind:side | 126 | 0.088 | 0.0006 | 0.018 | 1.0974 | 1.9296 | 0.0277 |
| Ind: side: replicate | 254 | 0.1617 | 0.0006 | 0.033 |  |  |  |
| Total | 507 | 4.8186 |  |  |  |  |  |
| **Plastron** |  |  |  |  |  |  |  |
| Ind | 126 | 11.451 | 0.0908 | 0.89707 | 58.522 | 23.1057 | <0.0001 |
| Side | 1 | 0.0183 | 0.0183 | 0.00144 | 11.8127 | 3.6142 | <0.0001 |
| Ind: side | 126 | 0.1957 | 0.0015 | 0.01533 | 0.3586 | -14.8321 | 1 |
| Ind: side: replicate | 254 | 1.0999 | 0.0043 | 0.08616 |  |  |  |
| Total | 507 | 12.765 |  |  |  |  |  |

Across species and ecological groups, asymmetry in the total shell configuration was generally concentrated in the anterior-left portion of the carapace, accompanied by posterior deformation of the shell (Figure 5A). In many species, the carapace showed pronounced asymmetry along the anterior margin, where left–right divergence occurred as localized widening on one side and narrowing on the other (Figure 5B). In contrast, plastral asymmetry was largely restricted to the bridge and hinge regions, with deformation patterns indicating unequal displacement of the anterior and posterior plastral lobes (Figure 5C).

**Figure 5.**
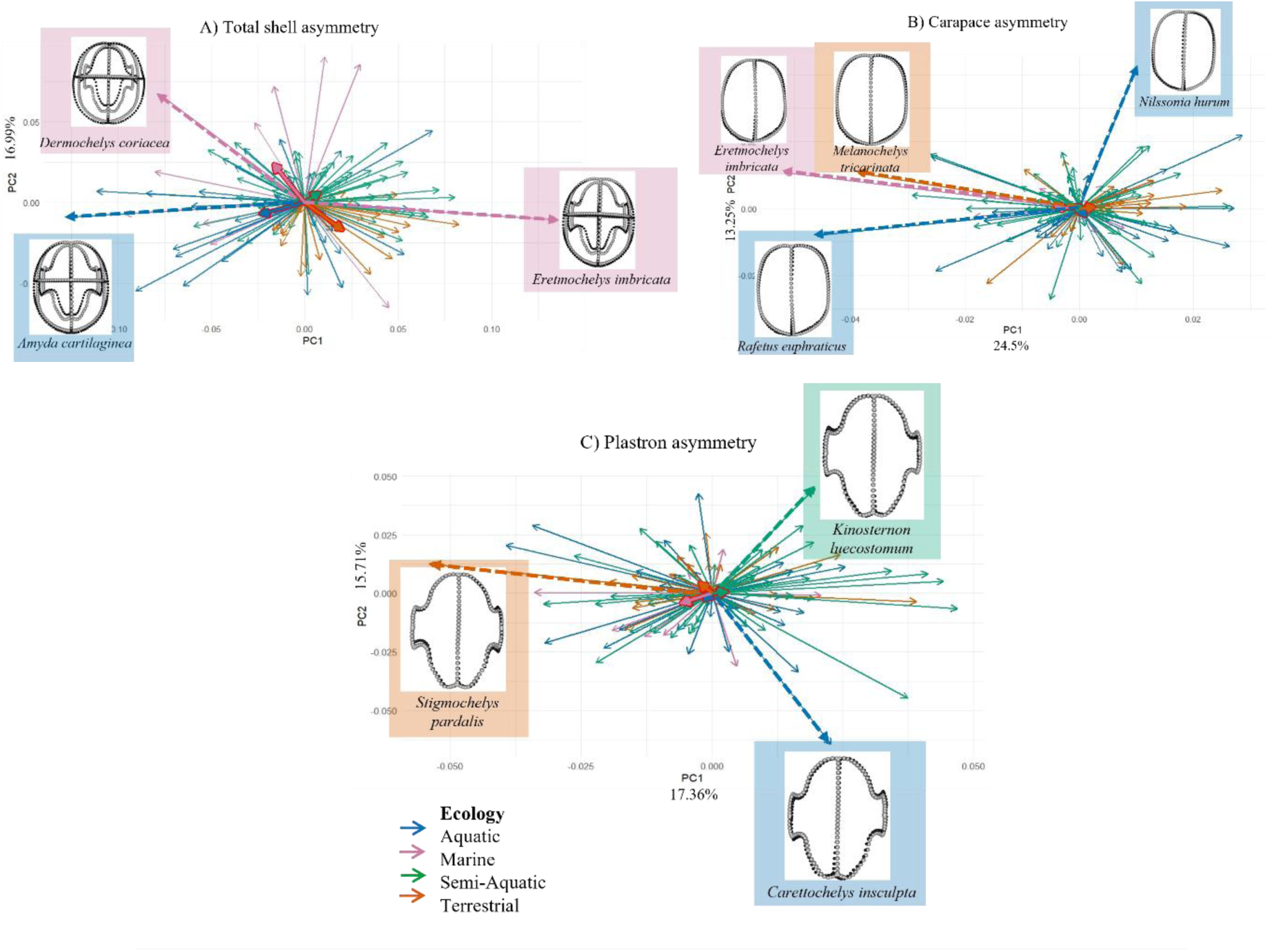
Principal component asymmetry scores of individuals, shown as arrows representing the magnitude of deviation from bilateral symmetry for the A) total shell, B) carapace, and C) plastron. Dashed arrow represents specimens having extreme shape asymmetry with respect to their ecology. Thick small arrows with red borders represent mean shape of each ecology group.

The degree of asymmetry varied significantly among ecological groups for the whole shell and plastron, but not for the carapace (Figure 6, Appendix 1). Among whole-shell measurements, the marine species *Lepidochelys kempii* exhibited the highest asymmetry, followed by the aquatic species *Chelydra serpentina* and *Chelydra acustrirosis*. For the carapace, higher asymmetry was observed in aquatic species such as *Rafetus euphraticus* and *Nilssonia hurum*, while the marine species *Eretmochelys imbricata* showed the highest value within its group. Notably, plastron asymmetry was substantially lower than both whole-shell and carapace asymmetry across all ecological categories, with *Batagur kachuga* and *Carettochelys insculpta* displaying the highest unsigned asymmetry values. See Appendix 3 for list of individual specimen AI values.

**Figure 6.**
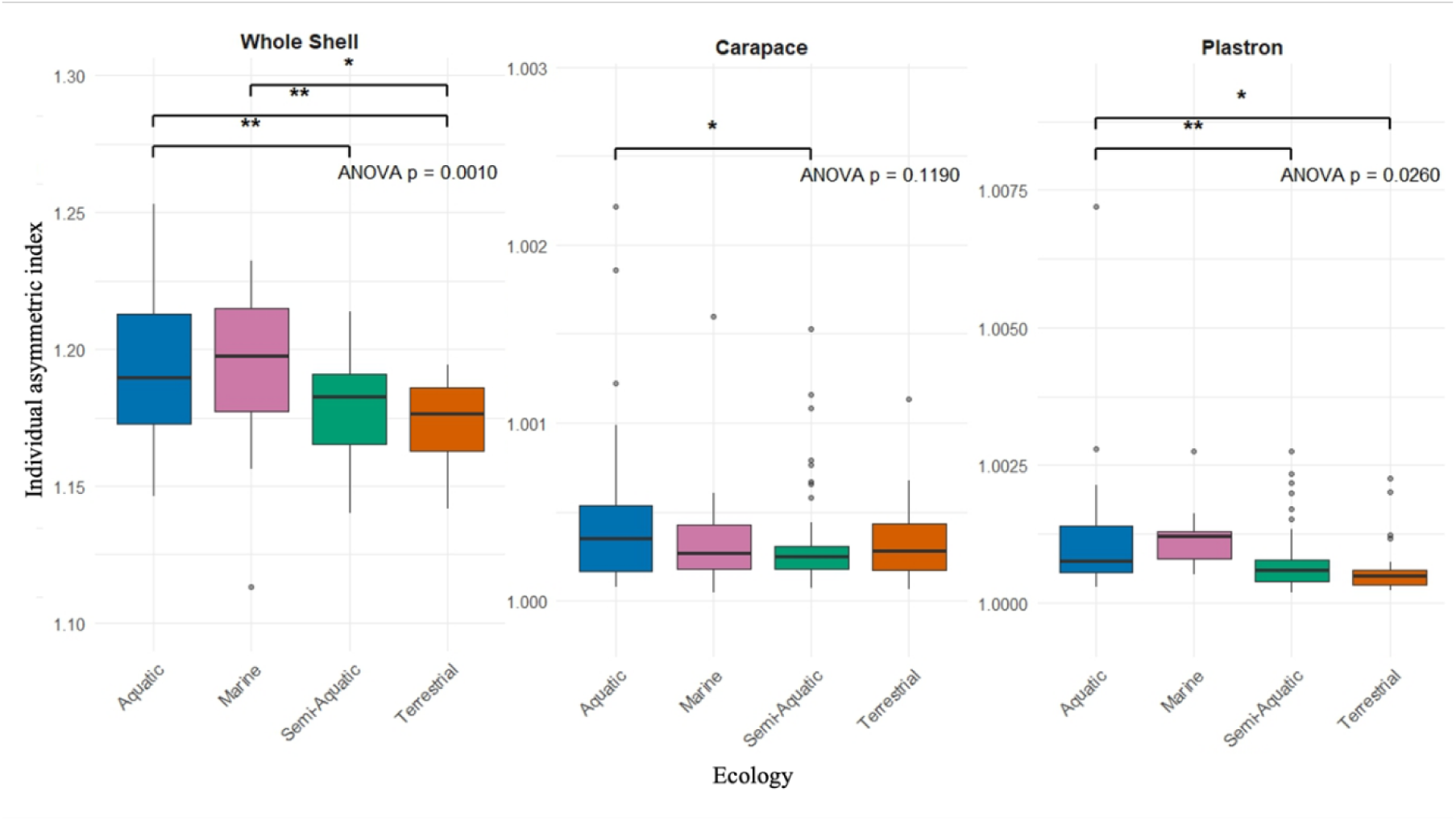
Box plots of turtle shell unsigned asymmetry index by ecology, representing the magnitude of deviation from bilateral symmetry in A) whole shell, B) carapace, and C) plastron. Asterisks indicate significant differences in asymmetry between ecological groups (* p < 0.05, *** p < 0.01). Note the differing y-axes.

The PLS analysis revealed significant covariation among symmetric and asymmetric patterns in the carapace and plastron (Figure 7). For the carapace (Figure 7A), symmetric and asymmetric components showed moderate integration, while the plastron exhibited a similar pattern (Figure 7B). Covariation between carapace and plastron symmetric components was stronger (Figure 7C), with a significant linear relationship also observed between their asymmetric components (Figure 7D). These results indicate that shape components within and between dorsal and ventral structures are not independent.

**Figure 7.**
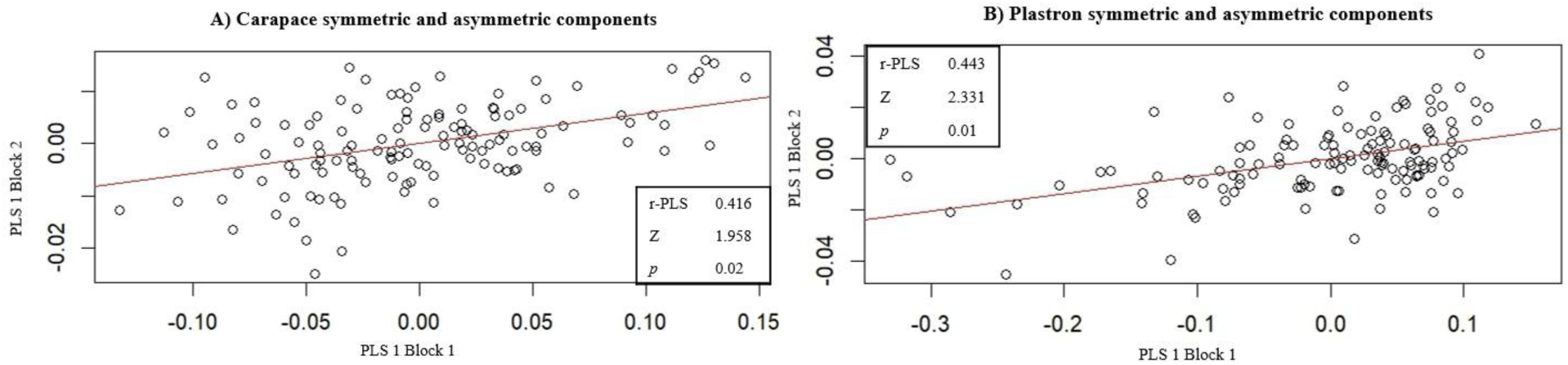
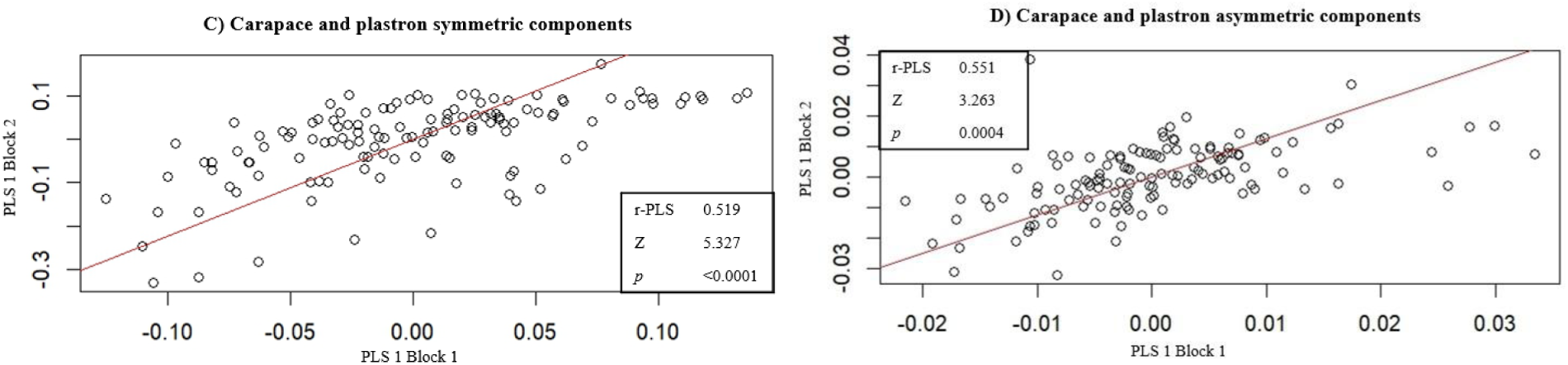
Result of two-block partial least square (PLS) analysis for carapace and plastron symmetric and asymmetric shape components, showing significant associations among all datasets.

## Discussion

Previous studies of 3D turtle shell shape used Type I (anatomical points) and Type II (curvature points) landmarks at the intersections of dermal or vertebral scutes [75–77]. In this study, we followed the approach of Dziomber et al. [6] that applied Type II and III landmarks and semi-landmark curves, presenting an opportunity to explore asymmetry by comparing left and right edges of the turtle shell [75]. This landmark scheme also allowed us to capture the dome shape of the carapace (Figure 2; C3 & C12) and curved midline and cross-sections of the plastron (Figure 2; C10 & C11), providing an overview of shell shape for phylogenetic and ecological comparisons. Using phylogenetic comparative methods, we detected a significant effect of ecology on shell shape, which may underlie observed patterns of directional and fluctuating asymmetry.

### Ecology is a strong factor in shaping turtle shell evolution

The scaling of biological traits with size, or allometry, is shown to be one of the strongest drivers of morphological variation within and between species [78]. Here evidence for allometry in Testudines was limited, as we sampled only adult specimens of varying sizes (except for one juvenile *D. coriacea*, see [6]), with sea turtles being the largest. We found a weak but significant effect of size on shell shape, which remained even after phylogenetic correction. Most turtles in our dataset are considered medium-sized, with the smallest centroid size estimated for the red-eared slider, *T. scripta*. Our addition of the remaining three living marine turtle species to the Dziomber et al. [6] dataset (*L. kempii*, *L. olivacea* and *N. depressus*), as well as the apparent ecological signal in centroid size, likely explains the lack of allometric signal found in their analysis (not incorporating phylogeny). Other studies such as Casale et al. [42] and Hermanson & Evers [7] recovered allometric relationships among turtles, indicating that the result is highly dependent on sample, number, type and location of landmarks, and application of phylogenetic comparative methods. We also found that evolutionary allometry varied among ecological groups, contrasting to previous findings where aquatic and terrestrial turtles belonging to Emydidae and Testudinoidea, respectively, showed similar allometric trajectories [76].

We found a significant effect of ecology on shell shape in both Procrustes and phylogenetic ANOVA analyses. This supports the idea that ecological factors play a key role in shaping morphological variation among turtles with different locomotory habits. Interestingly, the inclusion of phylogeny only slightly decreases this signal, suggesting that ecological adaptations are relatively independent of shared ancestry, further predicting the convergence of shapes in similar habitats [44,79,80]. This differs from ecomorphological patterns recovered for emydids (Emydidae, commonly known as pond or marsh turtles), in which ecological diversification tends to follow phylogenetic branching patterns [81,82]. However, it should be noted that the majority of turtle families have relatively conservative ecologies, meaning that their members belong to the same ecological group (i.e., terrestrial Testudinidae or aquatic Trionychidae, but see more ecologically diverse families such as Geomydidae; Figure 3).

In line with this pattern, the observed phylogenetic signal across the first twenty principal components (accounting for 95% of shape variation) reveals a significant but relatively weak morphological resemblance among closely related species, as indicated by the low K_*mult*_ value. However, the elevated traceK and detK values suggest that certain dimensions of shell shape retain stronger phylogenetic structuring, implying that evolutionary history still plays a role in shaping phenotypic traits. This conclusion contrasts with Mitteroecker et al. [66], who reported a greater sensitivity of phylogenetic signal metrics to dimensionality reduction, particularly due to the influence of small eigenvalues on K_*G*_. Our results show robust effect sizes and consistent significance across all metrics, suggesting that shell shape evolution in Testudines may be influenced by both phylogenetic history and ecological factors. Specifically, the stronger signal in plastron shape could reflect convergent adaptations to similar habitats, as previously reported [82]. For example, marine and soft-shell turtles tend to have reduced plastrons to minimize drag and improve swimming efficiency, while terrestrial tortoises have plastrons that are flat in females, with the males’ being often concave to facilitate mating [83,84]. Other terrestrial species, like box turtles, have a hinged plastron that allows them to close their shell completely, thus protecting their inner organs [85].

Apart from aquatic families Trionychidae and Chelydridae which formed discrete clusters along the first two principal components, many branches crisscrossed in phylomorphospace (Figure 4). This pattern suggests that shared phenotypic traits among distantly related species may occur in similar ecological niches, also known as convergent evolution. Studies comparing aquatic and terrestrial turtle clades have shown convergent flattening or doming of the carapace and plastron shapes associated with habitat transitions using different landmarking schemes [76,86]. Our pairwise ecological comparisons (Table 3) indicate a gradient of shell shapes that correlates with time spent in water, where fully aquatic turtles exhibit a distinct morphology that progressively differs as species become more terrestrial. These ecomorphological correlations align with previous research documenting how hydrodynamic constraints impose different selective pressures on shell shape compared to the mechanical demands of terrestrial locomotion [77,87,88], providing a mechanistic explanation for convergent turtle phenotypes.

### Significant directional and fluctuating asymmetry in turtle shells

Our Procrustes analyses partitioned morphological variation in turtle shells across 127 individuals from 92 species into components attributable to individual (species) differences, side effects (left-right asymmetry of whole shell, carapace and plastron configurations), and their interactions (fluctuating asymmetry). For the whole shell, the side effect explained 83.5% of total shape variation, vastly exceeding variation among species and fluctuating asymmetry. This dominant side effect reflects the shape differences between the articulated left and right dorsal (carapace) and ventral (plastron) shell components, which may differentially be shaped by ecological selection pressures. As illustrated in the arrow plot (Figure 5A), aquatic species possess highly flattened carapaces with expanded plastrons (blue vectors, *Amyda cartilaginea*), while terrestrial species exhibit domed carapaces and robust plastrons (orange vectors). These ecological specializations may underlie observed differences in left versus right morphology, where even slight rotations of the carapace relative to the plastron could result in large-scale side effects.

Directional asymmetry was negligible but significant in both the carapace (0.1%) and plastron (0.14%), indicating subtle but consistent left-right differences across turtle species. This low-level DA may reflect functional asymmetries related to locomotion, as suggested by previous studies in *Trachemys scripta elegans* where turtles showed preferential limb use [38]. Fluctuating asymmetry was significant only in the carapace (1.8%), but not in the plastron, suggesting that the plastron experiences stronger developmental canalization or stabilizing selection [89]. The carapace, being more exposed to environmental stressors and requiring greater structural integration with the vertebral column and ribs, is likely more susceptible to developmental perturbations [90]. As such even minor carapace asymmetry may impair reproductive performance and ultimately reduce individual fitness [10]. Other studies observed significant DA but no FA in *Testudo hermanni boettgeri*, suggesting that both the carapace and plastron grow consistently following their respective directional ossification patterns [91]. In *Gopherus polyphemus,* FA was detected to be highest in the population with the lowest genetic diversity [38], which should be further studied with other possible environmental stressors. Furthermore, DA and FA were significant in *Trachemys scripta elegans* plastron shape [38], suggesting that turtles might have preferential use of a thoracic or pelvic side for locomotion.

We observed habitat-specific influences on whole shell and carapace for aquatic and marine species, displaying higher individual asymmetric index values than terrestrial species. The reduction in plastron asymmetry compared to the carapace indicates differential stabilizing selection or developmental constraints between dorsal and ventral shell components across ecological niches. Individual variation in carapace and plastron shape was also highly significant and explains the majority of phenotypic diversity (94.6% and 89.7%, respectively), similar to the individual effects found in the head shape of lizards [16]. Sexual dimorphism was not accounted for in our study due to the lack of labeling on museum specimens. However, directionality can also be indicative of mating fitness, as observed in phallostethid fishes, where right-sided males exhibit greater fitness and left-sided females demonstrate higher dominance [92]. While similar effects have not been reported in turtles, greater DA has been observed in the carapace of males *Testudo* tortoises, which may prefer to roll on one side when self-righting after fighting with other males [37].

Measurement error was low in the whole shell and carapace, whereas the plastron showed higher values. This may be due to greater shape variation in plastrons across different species, especially around the bridge that connects the carapace—potentially making accurate landmark placement there difficult. Trionychid (softshell) turtles present a unique challenge in morphometric analyses due to their highly specialized plastron morphology. Unlike other testudines, softshell turtles lack clearly defined scutes and possess a reduced ossification pattern, making landmark identification and bridge shape determination particularly challenging [93,94]. This distinctive morphology represents an evolutionary adaptation to their highly aquatic lifestyle, reflecting their bottom-dwelling behavior and enhanced swimming performance [95,96]. The reduced plastron of trionychids, characterized by peripheral fontanelles and modified bridge connections, exemplifies how selective pressure drives morphological divergence in response to specific ecological demands [97,98]. In other studies, a strong symmetry signal was observed in aquatic species with low symmetry in terrestrial turtles belonging to the Testuguria superfamily, which includes the Geoemydidae and Testudinidae [77]. In contrast, terrestrial members of Emydidae (not sampled in our study) were found to be more symmetric than aquatic ones, suggesting that the inconsistency of asymmetry patterns among ecotypes is likely due to epigenetic or environmental factors rather than natural selection.

The PLS analysis revealed significant covariation between symmetric and asymmetric components of shell shape, with stronger integration observed in the plastron compared to the carapace. This suggests that the plastron may be more developmentally constrained or influenced by shared environmental factors during ontogeny. Interestingly, while the carapace showed moderate integration, its higher correlation and effect size may reflect a more complex developmental trajectory or delayed morphological integration. These findings align with a study on the evolution of hinge joints, which demonstrated that shell kinesis in the plastron arises repeatedly across distantly related turtle lineages, indicating a high degree of developmental plasticity and functional convergence [99]. Moreover, the coordinated covariation between dorsal and ventral shell components supports the hypothesis that the turtle shell functions as an integrated morphological unit, despite differences in developmental timing. Similar integration patterns have been observed in other reptiles, such as lizards, where anterior and posterior head regions show significant covariation across urban and rural populations [74], suggesting that environmental pressures can shape asymmetric traits in a coordinated manner. Overall, these results highlight the interplay between developmental constraints and ecological adaptation in shaping the evolution of complex morphological structures like the turtle shell. In the future, FA should be integrated with biomechanical models like Finite Element Analysis to estimate specific stresses and strains on asymmetrical regions [8], further illuminating potential fitness consequences of this phenomenon.

### Limitations of museum collections for detection of FA

While museum collections offer valuable access to diverse taxa across time and space, they can also limit studies of shell shape in turtle specimens. For example, shells that were wet (i.e., preserved in ethanol) or wax-coated were difficult to scan using the Artec Space Spider, due to light reflection off the surface. This could be mitigated by employing other 3D scanning techniques like X-ray CT, which offer better fidelity for complex surfaces yet fail to capture colour, which may be useful in other studies. Some surface scanners recommend spraying reflective specimens with a 3D scanning spray; however we found the powder spray difficult to fully remove after scanning. The absence of standardized labelling protocols in older collections like ZMUC also poses challenges, since missing metadata such as sex, locality, and collection year restricted our ability to reconstruct key biological and environmental variables. This is a common issue in museum-based research; over 90% of reptile specimens in the Western Australian Museum were found to lack sufficient geographic data for ecological studies [99]. Other specimens were excluded due to missing scutes or plastrons, further reducing our sample size. Finally, our landmarks focused on the shell edges and midlines (Figure 2), thus missing large, often curved surfaces of the carapace and plastron. Emerging methods like automated landmarking and 3D surface mapping could allow efficient placement of hundreds of landmarks across the entire shell surface, also improving reproducibility (i.e., reducing measurement error) in future studies [100].

### Conclusion: FA may be a sensitive tool to for monitoring health in living turtles

Our study demonstrates the utility of 3D surface scanning combined with geometric morphometrics and asymmetry analyses for quantifying shell shape variation in turtles. These non-invasive methods can be applied to living individuals across ontogenetic stages, species, and ecological communities to monitor developmental stability and detect early signs of stress. For example, fluctuating asymmetry has been linked to environmental disturbances in other taxa [16,19] and could serve as a sensitive indicator of habitat quality or physiological stress in turtles. Applying these approaches in wild populations would enable long-term monitoring of health and growth patterns under changing environmental conditions, such as pollution, temperature shifts, and habitat fragmentation.

Given the ecological roles of turtles as seed dispersers, scavengers, and regulators of diverse environments, maintaining their health is critical not only for species survival but also for ecosystem functioning. Future research should explore FA and shape quantification as tools for conservation, particularly in threatened habitats or populations exposed to anthropogenic pressures. Integrating morphometric monitoring with ecological and genetic data could provide a powerful framework for assessing population resilience and informing management strategies.

## Supporting information

Appendix 1: Classifiers table

Appendix 2: Centroid size table

Appendix 3: Unsigned asymmetry values table

Appendix 4: Landmark file for all individuals

Appendix 5: Landmark pairs file for whole shell

Appendix 6: Ecology file

Appendix 7: Landmark file for carapace

Appendix 8: Landmark pairs for carapace

Appendix 9: Landmark file for plastron

Appendix 10: Landmark pairs for plastron

Appendix 11: Landmark file for species averages

Tree file

## Acknowledgement

We gratefully acknowledge the Bio Research Advancement Fund at the University of Copenhagen for supporting this study. We thank Tristan Stayton for sharing CT data used to generate four shell models included in this work, Mark Scherz and Daniel Klingberg Johansson from the Natural History Museum of Denmark for providing access to collections and Kasper Lykke Hansen for assistance in surface scanning, and Marc Jones and Tom Ransom at the Natural History Museum of London for scanning the *L. kempii* specimen. Finally, we thank Barbara Fischer and Anne Le Maître for their valuable comments on the manuscript.

