## Supplementary document for "3D shell asymmetry of Testudines as a potential biomarker for environmental stress"

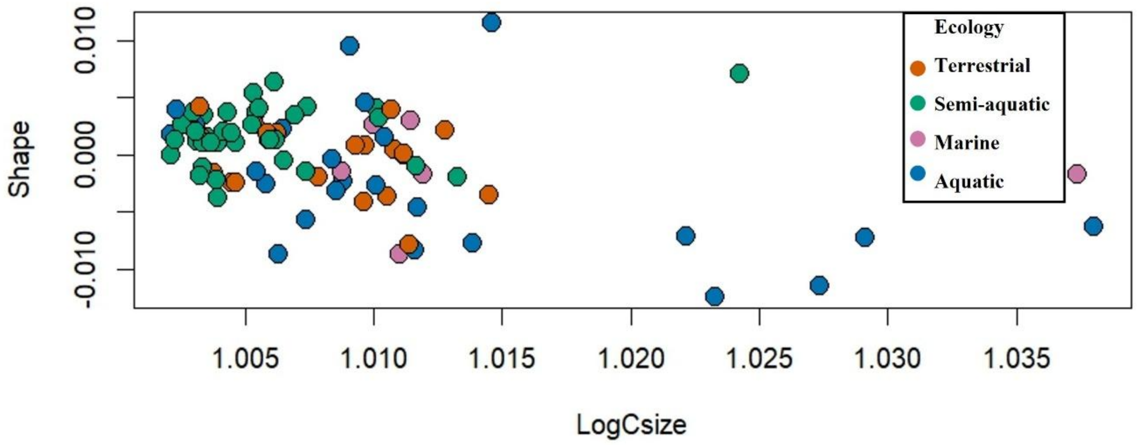


**Figure S1.** Allometric relationship between size and shell shape across 92 turtle species, sho wing PC1 fitted values plotted against the size predictor.

**Table S1.** Results of allometric analysis testing the relationship between centroid size (log Cs ize) and shape variation for all individuals.

| | Df | SS | MS | Rsq | F | Z | Pr(>F) |
|---|---|---|---|---|---|---|---|
| **log(Csize)** | 1 | 0.117 | 0.117 | 0.024 | 5.857 | 3.671 | <0.0001 |
| **Residuals** | 232 | 4.669 | 0.02 | 0.975 | | | |
| **Total** | 233 | 4.787 | | | | | |

**Table S2.** ANOVA testing the effect of ecological category on asymmetric shape variation of the whole shell.

| | Df | SS | MS | Rsq | F | Z | Pr(>F) |
|---|---|---|---|---|---|---|---|
| **Ecology** | 3 | 0.07447 | 0.0248225 | 0.12206 | 5.7464 | 4.3006 | 0.001 |
| **Residuals** | 124 | 0.53564 | 0.0043196 | 0.87794 | | | |
| **Total** | 127 | 0.6101 | | | | | |

**Table S3.** ANOVA testing the effect of ecological category on asymmetric shape variation of the carapace

| | Df | SS | MS | Rsq | F | Z | Pr(>F) |
|---|---|---|---|---|---|---|---|
| **Ecology** | 3 | 0.002451 | 0.000817 | 0.02765 | 1.1754 | 0.6816 | 0.242 |
| **Residuals** | 124 | 0.08617 | 0.000695 | 0.97235 | | | |
| **Total** | 127 | 0.08862 | | | | | |

**Table S4.** ANOVA testing the effect of ecological category on asymmetric shape variation of the plastron

| | Df | SS | MS | Rsq | F | Z | Pr(>F) |
|---|---|---|---|---|---|---|---|
| **Ecology** | 3 | 0.009257 | 0.003086 | 0.03049 | 1.2997 | 0.89203 | 0.186 |
| **Residuals** | 124 | 0.2944 | 0.002374 | 0.96951 | | | |
| **Total** | 127 | 0.30366 | | | | | |

**Table S5.** Pairwise comparisons of turtle shell shapes among ecological groups with phyloge netic correction.

| **Ecological comparison** | **r** | **Upper 95% CL** | **Z** | ***p*** |
|---|---|---|---|---|
| Aquatic vs. Marine | -0.542 | 1.285 | 3.388 | 0.0001 |
| Aquatic vs. Semi-aquatic | 0.556 | 1.263 | 0.742 | 0.228 |
| Aquatic vs. Terrestrial | -0.426 | 1.595 | 2.583 | 0.002 |
| Marine vs. Semi-aquatic | -0.397 | 1.457 | 2.816 | 0.002 |
| Marine vs. Terrestrial | 0.302 | 1.689 | 0.421 | 0.341 |
| Semi-Aquatic vs. Terrestrial | -0.026 | 1.687 | 1.425 | 0.084 |

**Table S6.** Results of ANOVA testing the effect of ecological category on unsigned shell shape variation.

A) Total shell shape

| | Df | SS | MS | Rsq | F | Z | Pr(>F) |
|---|---|---|---|---|---|---|---|
| **Ecology** | 3 | 0.007642 | 0.002547 | 0.12219 | 5.7069 | 2.93525.2337 | 0.001 |
| **Residuals** | 123 | 0.0549 | 0.000446 | 0.87781 | | | |
| **Total** | 126 | 0.06255 | | | | | |

B) Carapace

| | Df | SS | MS | Rsq | F | Z | Pr(>F) |
|---|---|---|---|---|---|---|---|
| **Ecology** | 3 | $7.0260 \times 10^{-7}$ | $0.02.3419\times 10^{-7}$ | 0.04514 | 1.9384 | 1.1359 | 0.119 |
| **Residuals** | 123 | $1.486 \times 10^{-5}$ | $1.2081\times 10^{-7}$ | 0.95486 | | | |
| **Total** | 126 | $1.5563\times 10^{-5}$ | | | | | |

C) Plastron

| | Df | SS | MS | Rsq | F | Z | Pr(>F) |
|---|---|---|---|---|---|---|---|
| **Ecology** | 3 | $6.5990 \times 10^{-7}$ | $2.1996\times 10^{-6}$ | 0.07937 | 3.53451.9384 | 2.1154 | 0.026 |
| **Residuals** | 123 | $7.6545 \times 10^{-5}$ | $6.2231\times 10^{-7}$ | 0.92063 | | | |
| **Total** | 126 | $8.3143\times 10^{-5}$ | | | | | |
